# From exposure to infection: divergent fitness consequences of parasite encounters in a trophically-transmitted system

**DOI:** 10.64898/2026.05.06.723225

**Authors:** Chloe A. Fouilloux, Jonathan S. Compton, Ipsita Srinivas, Marisa L. Schuldes, Audrey L. Rollo, Ryan Paulman, Josh Sampson, Amanda Hund, Jessica L. Hite

## Abstract

Parasites can alter host populations in fundamentally different ways depending on whether exposure results in infection. Yet, most epidemiological and evolutionary inference focuses on established infections, leaving the fitness consequences of parasite exposure comparatively understudied. This gap is consequential because hosts are frequently exposed to diverse parasite genotypes, and these encounters can impose substantial fitness costs even when infection does not occur. Theory predicts that hosts may mitigate these costs when interacting with commonly encountered parasite genotypes, such that exposure to sympatric parasites incurs lower fitness consequences than exposure to novel, allopatric parasites. Here, we examine the fitness consequences of exposure and infection in the first intermediate host of the trophically transmitted tapeworm *Schistocephalus solidus*, a cyclopoid copepod that serves as the first host in a three-host life cycle. Using sympatric (Vancouver Island, Canada) and allopatric (Norway) host-parasite combinations, we found a striking reciprocal asymmetry. Sympatric parasites were significantly more infective, yet exposure to sympatric parasites imposed weaker fitness costs when infection did not establish. In contrast, allopatric parasites were less infective, but exposed females produced fewer eggs and had lower hatching success than both controls and females exposed to sympatric parasites, indicating substantial genotype-dependent costs of exposure. Moreover, we found that infection was highly virulent across all genotypes: a single parasite caused near-complete reproductive suppression and reduced host survival across all host-parasite pairings, confirming *S. solidus* as a castrating parasite in copepods. Together, these results demonstrate that exposure, not just infection, acts as a critical ecological filter with potentially large and underappreciated consequences for host population dynamics and parasite transmission.

## Introduction

Most host-parasite encounters end without infection, yet ecological and evolutionary inference is drawn almost exclusively from the small fraction of infections that succeed (Kuris et al. 2007). If most encounters terminate before infection, then focusing on only infected hosts captures a subset of the evolutionary and ecological pathways that drive host-parasite dynamics. Infections that do not establish can fail through several mechanisms including mismatch in host and parasite genotypes (Salvaudon et al. 2007, Zavodna et al. 2008), constitutive immune defenses (Armitage et al. 2003, Rolff and Siva-Jothy 2003) and induced immune responses following exposure (Sadd and Schmid-Hemple 2006, Kurtz 2007).

Each of these resistance mechanisms carries energetic costs that can affect host fitness when infection fails to establish. Avoiding infection can impose costs through immune activation and associated tissue damage (Lochmiller and Deerenberg 2000, Moret and Schmid-Hempel 2000, Hall et al. 2007). A small number of studies indicate the impact of these exposures can accumulate to measurable fitness costs (Pfenning-Butterworth et al. 2023). Despite this, these ‘trait-mediated’ effects remain poorly resolved across systems (Kuris 1980, Horn and Luong 2018, Gowler et al. 2023). Successful infections, on the other hand, consistently redirect host resources toward parasite growth, often severely reducing host reproduction or survival (Gandon et al. 2002, Hurd 2001, Bonds 2006). In extreme cases, parasites castrate hosts entirely, halting reproduction and altering population age and size structure (O’Keefe and Antonovics 2002, Bonds 2006, Best et al. 2010). Considering the full spectrum of encounter outcomes, from exposure to infection, is therefore necessary to understand how parasite encounters shape host fitness and population dynamics (Froelick et al. 2021, Goodnight et al. 2025).

The outcomes of parasite encounters are especially important in the first-intermediate hosts of parasites with complex life cycles. These hosts act as evolutionary gatekeepers: they determine which parasite genotypes reach later developmental stages and thus experience selection in definitive hosts (Froelick et al. 2021, Goodnight et al. 2025). Accumulated across a season, differences in survival, fecundity, and offspring viability in host populations can propagate upward through food webs, shaping both host population growth and the pool of susceptible hosts available to subsequent parasite generations (Wedekind 1997, Lafferty and Kuris 2002, 2009, Bass et al. 2021). Despite the critical position of first-intermediate hosts for disease transmission and foodweb stability, the ecological and evolutionary dynamics of these (often small invertebrate) hosts remain a black box (Blasco-Costa and Poulin 2017). In particular, we lack a clear understanding of how exposure affects host fitness and whether these costs vary among parasite genotypes. Such variation could fundamentally shape how host responses feedback to shape parasite transmission.

Theory and empirical studies offer competing predictions for how infectivity and the fitness costs of exposure should covary across parasite origins (Lively et al. 2004, Greischar and Koskella 2007, Gandon et al. 2008). Under local adaptation, sympatric parasites are expected to be highly infective but relatively cheap to resist due to coevolutionary matching (Carius et al. 2001, Refardt and Ebert 2007), whereas allopatric parasites may be less infective but impose greater fitness costs upon exposure if hosts are not adapted to these parasite genotypes. Alternatively, if fitness costs scale with parasite performance rather than parasite origin, more infective parasites may impose greater costs (Boots and Haraguchi 1999, Gandon and Michalakis 2000). Despite the importance of these predictions for understanding host-parasite dynamics, few empirical studies have jointly quantified parasite performance and the fitness consequences of exposure across parasite origins, particularly in first-intermediate hosts where most encounters are resolved.

Here, we test for genotype-specific asymmetries in infectivity and exposure costs using the trophically transmitted tapeworm *Schistocephalus solidus* and its cyclopoid copepod host as a case study. This three-host parasite must first infect a cyclopoid copepod before transmission to threespine stickleback fish and, ultimately, piscivorous birds (Wedekind 1997). The copepod stage represents the system’s earliest and most severe transmission bottleneck: free-swimming coracidia have limited time (under 36 hours) to locate and penetrate a host, and infected copepods experience reduced growth and fecundity (Wedekind 1997, Fig. 1). At this stage, the relative fitness consequences of successful infection versus exposure, and whether these consequences vary by parasite genotype, remain unresolved. To address these gaps, we exposed gravid female copepods (*Acanthocyclops robustus*) to sympatric (Vancouver Island, Canada) and allopatric (Norway) *S. solidus* genotypes and quantified both parasite infectivity and the fitness consequences of how exposure and infection alter fecundity, reproductive timing, hatching success, and survival of first-intermediate hosts.

**Figure 1.**
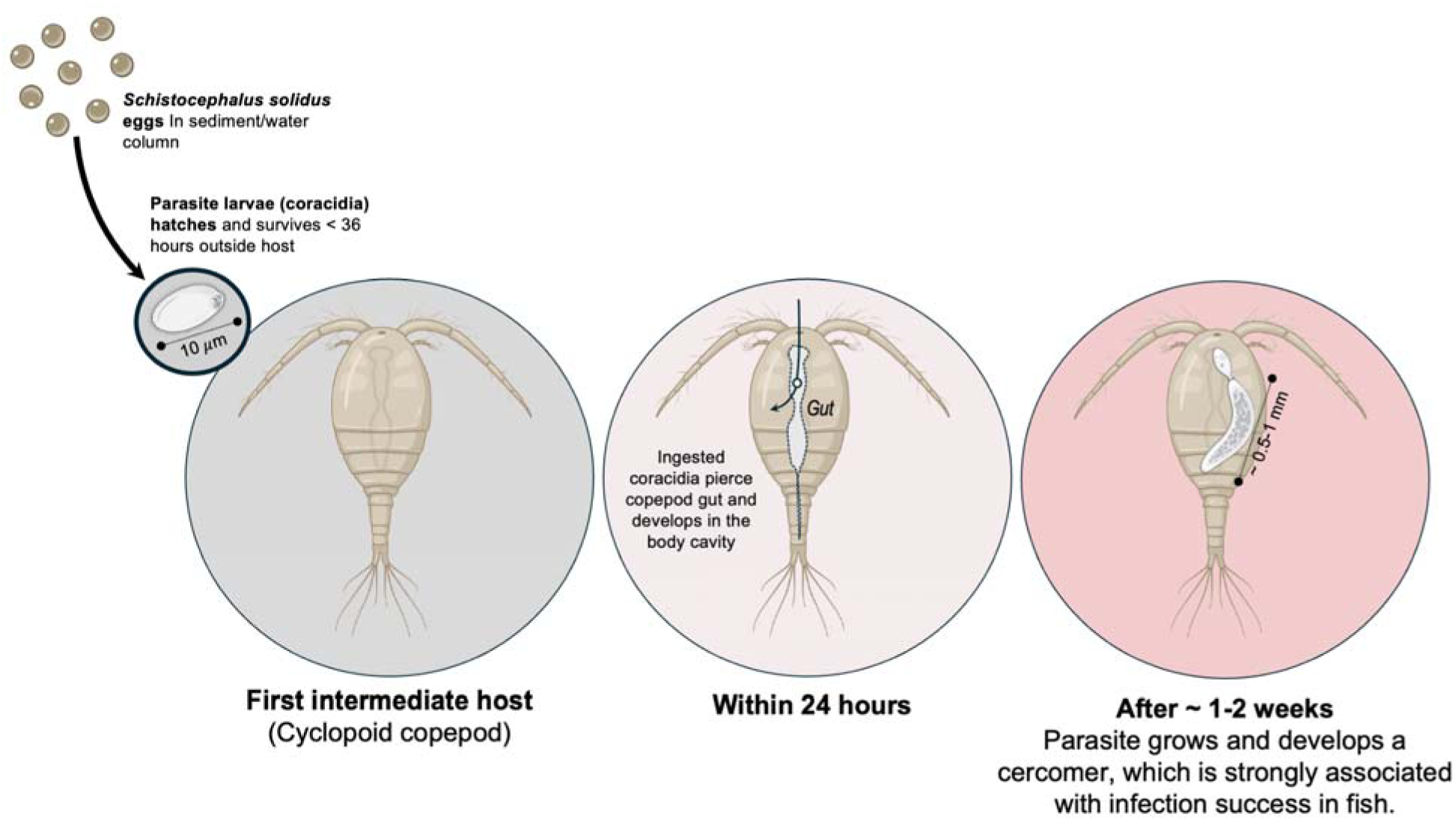
Overview of infection dynamics of *S. solidus* in copepod hosts. Parasite eggs hatch and larvae (now coracidia) must find a suitable host within 36 hours (left circle). Cyclopoid copepods ingest coracidia, which must successfully pierce the gut and enter host hemocoel (middle circle). Over the period of 1-2 weeks, procercoid parasites grow, developing structures (cercomer) associated with infection success in the subsequent intermediate host.

### BOX 1 Defining castration

While definitions and thresholds of parasitic castration vary across systems, here we adopt a functional, intensity-independent definition. Under this framework, castration is identified by a consistent parasite-associated suppression of host reproductive output relative to uninfected hosts, even at single-parasite intensity levels (Sorensen and Minchella 2001, Lafferty and Kuris 2002, Bass et al. 2021). The result of castration is reproductive death of the host, whereby the host becomes an extended phenotype of the parasite (Dawkins 1999, Lafferty and Kuris 2009).

## Methods

### S. solidus infection of its first intermediate host

Cued by light and temperature, *Schistocephalus solidus* eggs hatch into free-swimming larval parasites called coracidia, which must locate and infect a suitable host within approximately 36 hours (Wedekind 1997). Upon encounter, cyclopoid copepods ingest coracidia (Hammerschmidt and Kurtz 2009). Following ingestion, coracidia penetrate the gut epithelium and enter the hemocoel (Jakobsen et al. 2012), where they rapidly develop into procercoid larvae within hours (< 3 h; Hammerschmidt and Kurtz 2009). Over the subsequent one to two weeks post-establishment (mean = 9 days; Hammerschmidt and Kurtz 2009, Benesh and Hafer 2012, CAF obs.), procercoids develop a posterior structure called a cercomer. Although the precise function of the cercomer remains unresolved (Benesh 2010, Jakobsen et al. 2012), its presence is strongly associated with successful transmission to the second intermediate host, the threespine stickleback (*Gasterosteus aculeatus*) (Jakobsen et al. 2012). Cercomer development thus sets a minimum persistence time: *S. solidus* must survive in the copepod long enough to become infectious before killing it.

### Cyclopoid copepod system overview

Cyclopoid copepods reproduce sexually, and females typically mate once and subsequently produce sequential clutches from stored sperm that they continue to produce over a period that ranges from weeks to months (Maier 1995, Marten and Reid 2007). Mated females extrude eggs into lateral egg sacs, which they carry for a period of days (2.5-4.1 days; Hopp et al. 1997, Caramujo and Boavida 1999). Interclutch duration is remarkably consistent across successive clutches and does not vary substantially with clutch order in freshwater cyclopoids (Jamieson and Santer 2003), suggesting that the timing of clutch production is a relatively fixed physiological process. While the timing clutch production of cyclopoid females remains an area of ongoing research, the size of clutches relative to females is large (Fig. 3), and has been hypothesized to increase predation probability by fish (Maier 1995). This increased predation risk is thought to impose strong selection on reproductive scheduling and investment strategies in cyclopoid copepods (Maier 1995, Hopp et al. 1997).

### Stock culture maintenance and experimental individuals

To quantify how parasite exposure and infection alter the fitness of first-intermediate hosts, we conducted a life table assay using gravid adult female *A. robustus* copepods. This approach is tractable because female sperm storage follows single mating, and necessary given that female responses to infection remain largely uncharacterized in this system (Kurtz et al. 2002, Hammerschmidt et al. 2009, Benesh and Hafer 2012). These animals were selected from existing laboratory cultures isolated from Echo Lake (Vancouver Island, B.C., Canada) in 2015/2016 and maintained under standard laboratory conditions. More specifically, copepod hosts (∼ 600 adults L^-1^) were maintained in 1 L flasks at 19°C with a 16:8 L:D cycle with 900 mL of standard (low-hardness) COMBO water for animals (artificial lake water media, Kilham et al. 1998). Animal stocks were fed freeze-dried crushed *Artemia* (AMZEY Natural Artemia; 0.022 mg L^-1^) twice weekly. Flask populations were not mixed and maintained as separate lines.

To minimize variation in reproductive history among experimental individuals, we isolated gravid adult females bearing large egg clutches (mean clutch size = 56.3 eggs) from stock cultures, consistent with published values for early reproductive output in *A. robustus* (mean clutch size = 55.6 eggs; Maier 1995). Because the precise age and reproductive history of individual females prior to isolation were unknown, clutch number is denoted as (n+1) across visualizations. Survival of unexposed females in our study (mean lifespan = 56.8 days) closely matched published estimates for adult *A. robustus* under laboratory conditions (mean lifespan = 55.9 days; Hopp et al. 1997), indicating that experimental animals maintained normal physiological baselines.

### Experimental infection assays

We conducted dose-controlled exposures of two distinct *S. solidus* genotypes to assess host susceptibility and parasite infectivity across genotypes. Infected stickleback were originally collected from Kjerag Fjord, Norway (8 September 2022, GPS: 67.501487, 14.742647) and from Merrill Lake, Vancouver Island, Canada (2024, GPS: 50.060414, - 125.561481). In a laboratory setting, stickleback were dissected and *S. solidus* plerocercoids were collected and bred using previously described protocols (Smyth 1946, Wedekind 1997, Jakobsen et al. 2012, Weber et al. 2017). Briefly, following fish dissection, tapeworms from each lake were size-matched and paired. Tapeworm pairs were placed within nylon biopsy bags which were suspended in bottles filled with media (detailed in Jakobsen et al. 2012); breeding between tapeworms was stimulated by maintaining pairs in a shaking warm water bath (42°C) that simulated avian host environments.

Eggs were then harvested and stored. To minimize fungal growth, eggs were washed several times with sterile water (Weber et al. 2017) and maintained in long-term storage in the darkness at 4°C in 15mL falcon tubes at the University of Wisconsin, Madison. To stimulate egg hatching, 200µl of egg suspension was aliquoted into a single well of a foil-covered 24-well microtiter plate with 2mL of COMBO media and incubated in the dark at 18°C for seven days. Following this, egg plates were moved to room temperature (25-26°C) and placed under full-spectrum grow lights (GT-Lite LED Grow Bulb; 13.09 PPF, 8.5 Watts; SKU:GR-A19) with a 16:8 L:D cycle (Jakobsen et al. 2012, Weber et al. 2017). Hatching success was similar between both sympatric and allopatric parasite genotypes.

On the day prior to parasite exposure, single adult gravid female copepod hosts were isolated and in fresh COMBO and maintained without food in the 24-well plates with 1.5mL COMBO at 19°C under a 16:8 L:D cycle (Jakobsen et al. 2012). Following the starvation period, three *S. solidus* coracidia were placed in each well for a 48-hour exposure period. *S. solidus* coracidia survives less than 36 hours outside of its first intermediate host (Wedekind 1997). Given the combined 72-hour window of starvation and exposure, we expect all coracidia should be consumed by hosts (Hammerschmidt and Kurtz 2009). We verified this by examining the 24-well plates under a compound microscope (Leica M80 at 60x magnification using a Leica KL 300 LED) for any live or dead coracidia prior to transferring exposed gravid females to 15 mL falcon tubes. We did not detect any coracidia in exposed treatments.

The transparency of *A. robustus* allowed infections to be assessed *in vivo.* Infections were confirmed 9-14 days post-exposure (see Fig. 1) using a compound microscope (Leica M80 at 60x magnification using a Leica KL 300 LED). Infective intensity was confirmed post-mortem by dissecting tapeworms from exposed hosts. Experimental treatments consisted of unexposed females (n = 48), infected females (n = 61, where n_Local_ = 52 and n_Foreign_ = 9) and exposed-but-uninfected females (n = 58, where n_Local_ = 30 and n_Foreign_ = 28). By exposing a single host population to sympatric and allopatric parasites, we can directly test whether parasite performance and host fitness consequences reflect local adaptation. Because all *A. robustus* hosts originated from Vancouver Island, we henceforth refer to the Norwegian and Vancouver Island parasite genotypes as ‘allopatric’ and ‘sympatric’ respectively.

### Gravid female maintenance

To characterize the fitness consequences of exposure and infection, we tracked individual female reproductive output and survival daily until death. Immediately following the 48-hour exposure to *S. solidus*, individuals were moved to 15mL falcon tubes filled with COMBO and maintained at 19°C with a 16:8 L:D cycle. Tubes were filled with 0.4 to 0.67 particles/mL of *Artemia* suspension where females could feed *ad libitum.* Females were checked daily and water/food was changed weekly. Clutch size, clutch symmetry, intra/interclutch period, days to death post-isolation, total number of eggs post-isolation, and F1 survival were recorded.

## Statistical analysis

To partition the effects of treatment, clutch number, and individual identity on host life-history outcomes, we fit a series of mixed models. All analyses were conducted in R (v4.5.1, R Core Team 2025). Models were fit using `glmmTMB` (v1.1.12, Brooks et al. 2017) and evaluated using `DHARMa` (v0.4.7, Hartig 2024). Unless otherwise noted, count responses were modelled with a negative binomial distribution using the linear variance parameterization (`nbinom1`) and a log link. Model adequacy was assessed using simulation-based residual diagnostics; dispersion and zero inflation were modelled when supported by diagnostics. Because few hosts became infected following exposure to the allopatric parasite genotype (n = 9), infected outcomes were pooled across parasite genotypes. Differences in treatment group sizes were accommodated by likelihood-based estimation. As females were isolated gravid, models consider post-isolation egg and nauplii production to account only for treatment related effects.

Survival following isolation was analysed using Kaplan-Meier estimators and compared among treatments using log-rank tests and Cox proportional hazards models implemented in the `survival` package (v. 3.8.3, Therneau 2024).

### Egg counts in clutches

Clutch egg totals were modelled (m1) as a function of clutch number, inter-clutch interval, and their interaction with treatment (4-level factor: Unexposed, Exposed-Sympatric, Exposed-Allopatric, Infected), with random intercepts for female ID and stock. Residual variance and zero inflation were modelled at the intercept level.

### Lifetime egg and nauplii production

Post-isolation lifetime egg (m2) and nauplii totals (m3) (excluding clutch 1) were modelled as a function of treatment (4-level factor), with a random intercept for flask populations. For nauplii totals, zero inflation was modelled at the intercept level.

### Proportion of nauplii hatching from clutches

The proportion of nauplii hatching per clutch was modelled (m4) using a beta-binomial distribution with a logit link function, specified as successes and failures. Fixed effects included treatment and clutch number, with random intercepts for female ID and stock.

### Clutch timing models

Mean holding time (t1) and time between clutches (t2) were log-transformed and modelled as a function of treatment (4-level factor), with a random intercept for stock. For both responses, residual variance was allowed to vary across treatment when supported by model diagnostics. The oldest age at which a viable F1 was produced (days) (t3) was modelled as a function of treatment, with a random intercept for flask population, using a negative binomial distribution to accommodate overdispersed day counts. Total female lifespan (days survived post-isolation) (t4) was analysed using a log-transformed Gaussian mixed model with treatment as a fixed effect and stock as a random intercept.

## Results

### Sympatric genotypes exhibit higher infection success

To test for potential genotype-level differences in infection success, we exposed females to either sympatric or allopatric parasite genotype. Infection prevalence differed markedly between treatments (Fig. 2A). For sympatric parasites, the prevalence of infection was 63% (52 infected of 82 exposed). For allopatric parasites, the prevalence of infection was substantially lower, 24% (9 infected of 37 exposed). These notable differences indicate that parasites originating from the same host population were substantially more likely to establish infections relative to allopatric parasites.

**Figure 2.**
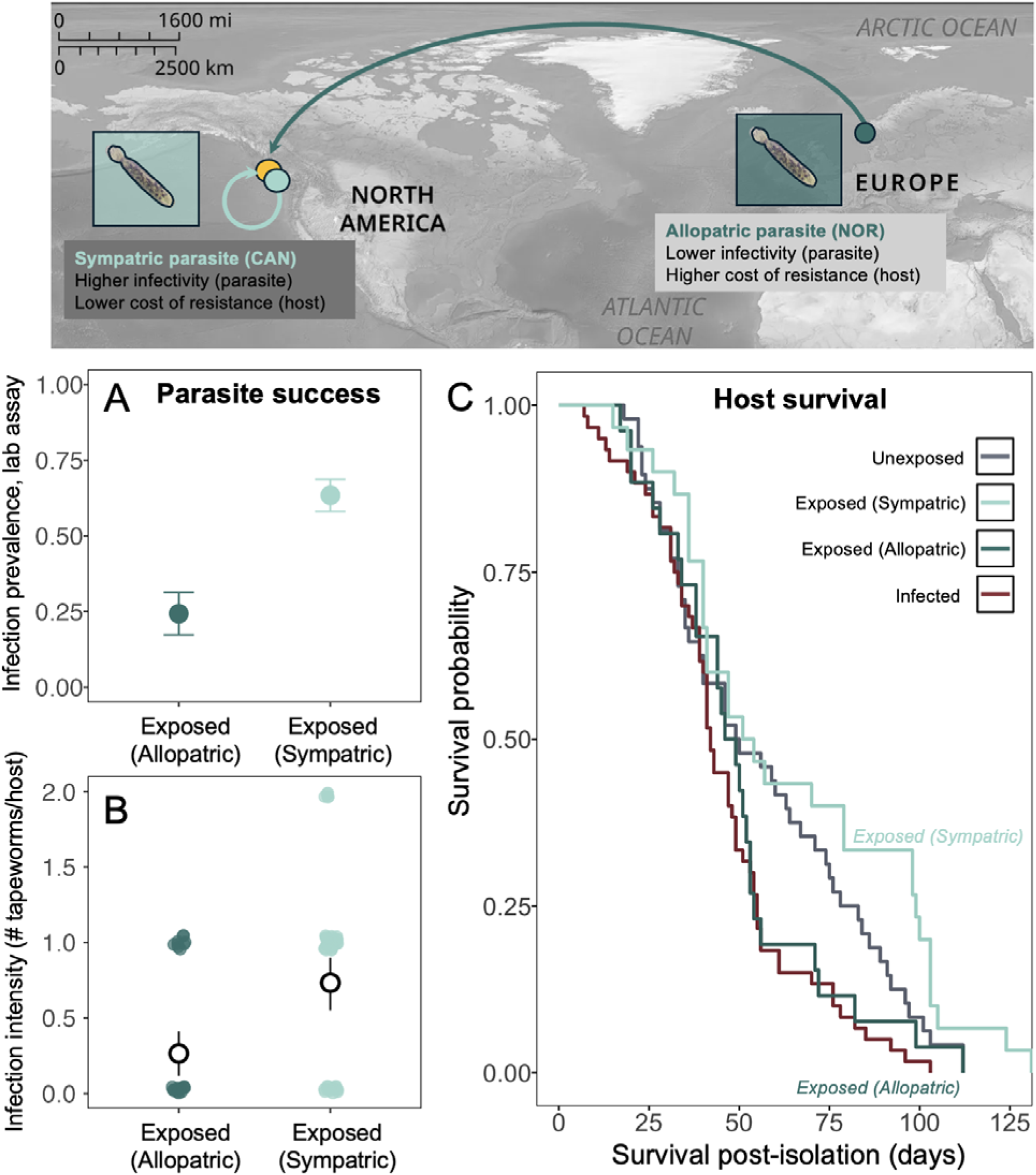
Reciprocal asymmetry between sympatric and allopatric host-parasite combinations. Sympatric parasites had higher infection success compared to allopatric parasites (A, B). However, females exposed but uninfected by sympatric parasites produced survived significantly longer post-isolation compared females exposed to allopatric parasites (C). Yellow point represents host origin (Echo Lake, Vancouver Island, Canada); sympatric parasites originate from Vancouver Island, Canada (Merrill Lake) and allopatric parasites originate from Norway (Kjerag Fjord).

Infection intensity also differed between parasite genotypes. Each host was exposed to three coracidia under controlled infection assays. All successful allopatric infections consisted of a single parasite per host (mean intensity among infected hosts = 1.0), yielding a mean intensity of 0.26 parasites across all exposed hosts. In contrast, sympatrically infected females harbored a mean of 1.29 parasites among successfully infected hosts (with ten individuals harboring two parasites), resulting in a mean intensity of 0.73 parasites per exposed host (Fig. 2B).

Expressed as per-coracidia establishment probability, 24% of sympatric parasites successfully established infections compared to 8.8% of allopatric parasites. Together, these results indicate that sympatric parasites were more likely to establish infections and achieve higher burdens within hosts. However, inference about differences in infection intensity should be interpreted cautiously given the limited number of allopatric infections.

### Host survival depended on infection outcome and parasite origin

To examine how exposure and infection affected host survival, we fit a Cox proportional hazards model across treatment groups. Treatment significantly affected mortality risk (likelihood ratio test: χ² = 13.9, df = 3, p = 0.003). Infected females exhibited significantly higher mortality than both unexposed controls (hazard ratio = 1.56, 95% CI = 1.06-2.3, p = 0.039; Fig. 2C), indicating strong survival costs associated with successful infection.

In contrast, exposure without infection had strong genotype-dependent effects on survival. Females exposed to allopatric parasites died significantly earlier than those exposed to sympatric parasites (hazard ratio = 1.88, 95% CI = 1.09-3.24, p = 0.022; Fig. 2C), whereas survival of sympatrically-exposed females did not differ from unexposed controls (95% CI = 0.422-1.08, p = 0.106). Indeed, the longest post-isolation lifespan recorded across all treatments was in a sympatrically-exposed female (131 days; ∼4.3 months), underscoring that sympatric exposure carried negligible survival costs when infection did not establish.

Taken together with infection outcomes, these results indicate that sympatric parasites are more likely to establish infections, but when infection is avoided, exposure to sympatric parasites is associated with lower survival costs relative to exposure to allopatric parasites.

### Fitness consequences resulting from tapeworm infection

Infection by *S. solidus* results in near-complete castration of female hosts. We quantified the fitness consequences of infection by tracking egg production and nauplii hatching across host lifespan. Infected females produced significantly fewer eggs (GLMM_m2_, z = -8.93, p < 0.001) and fewer nauplii (GLMM_m3_, z = -9.18, p < 0.001) than females in all other treatments, including unexposed controls (Fig. 3, Fig. 4B). These effects were consistent across parasite genotypes. None of the allopatrically infected females produced viable offspring (0/9), and only six of 52 sympatrically infected females produced viable nauplii, indicating that infection is associated with severe reductions in female reproductive output irrespective of parasite genotype (Fig. 4B).

**Figure 3.**
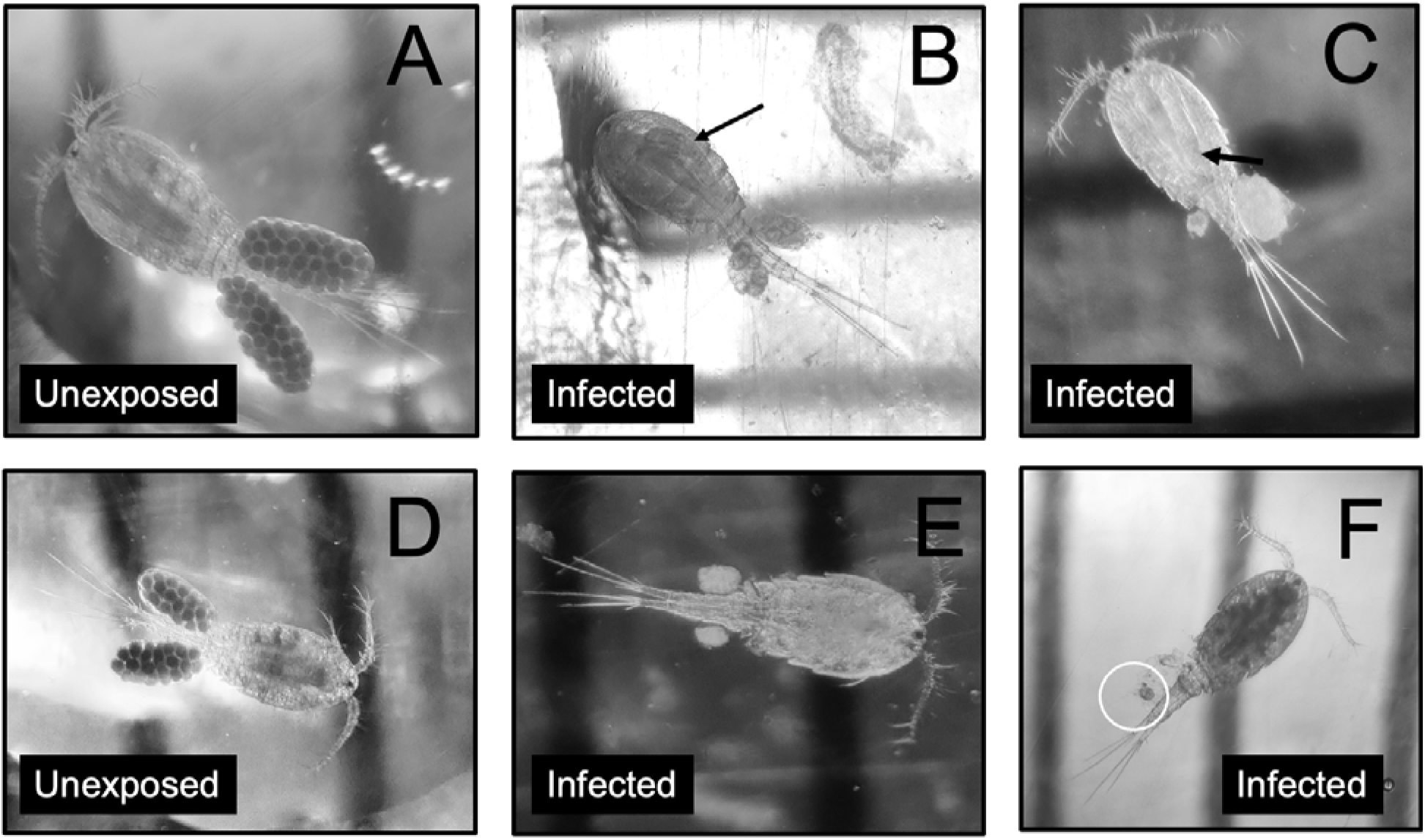
Quality of clutch development in infected females. Infected female photos represent common clutch phenotypes that failed to produce nauplii. Arrows in (B,C) highlight *S. solidus* tapeworm. Circle in (F) emphasizes a single-egg clutch. Lines in the image background represent 1mm scale.

**Figure 4.**
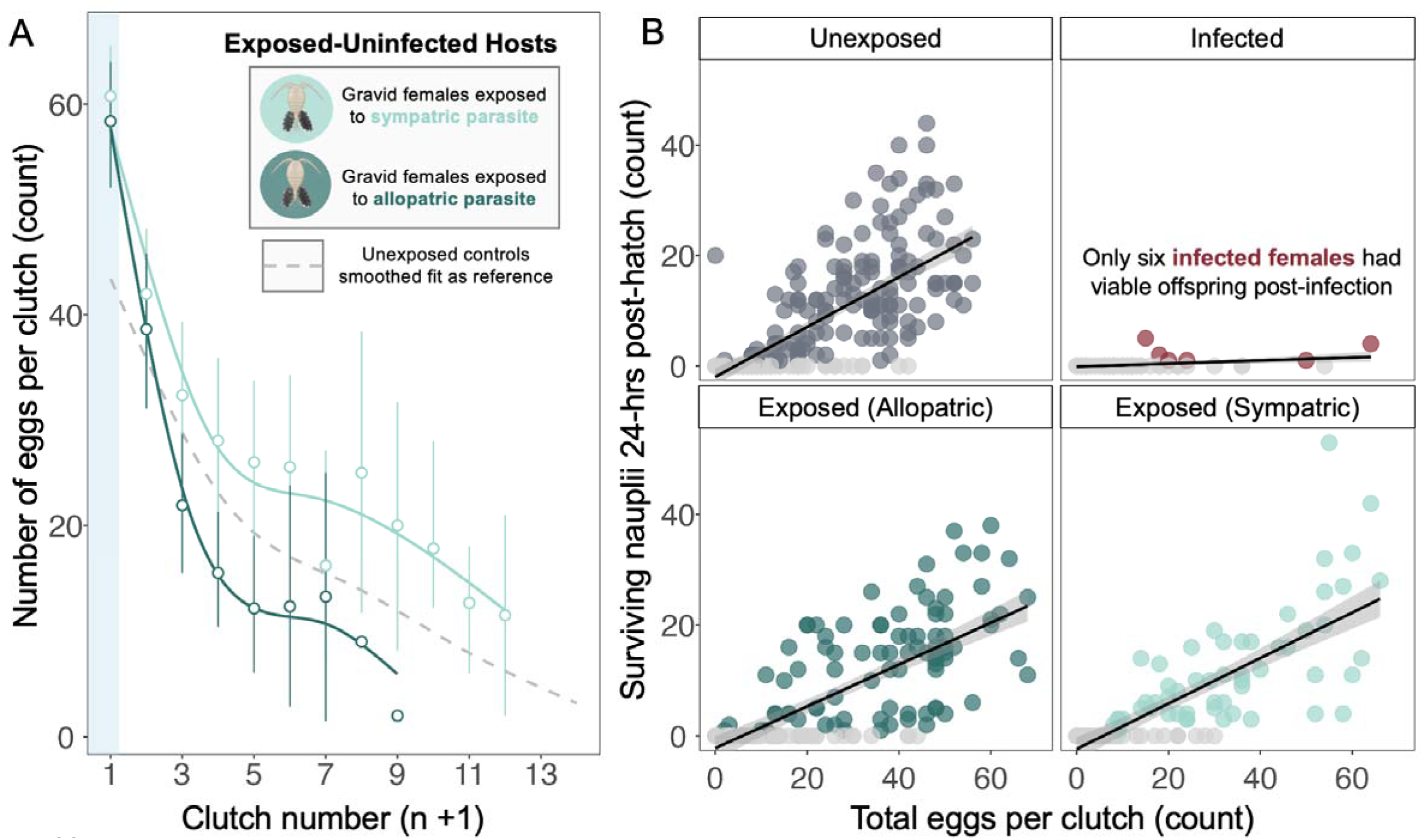
Genotype-specific costs of resistance. Females exposed to allopatric parasites produced significantly fewer eggs across clutches compared to those exposed to sympatric parasites (A). This translated to overall fitness outcomes, where proportion of viable nauplii was significantly lower in allopatric exposed females compared to controls (B). This is in contrast to near-complete reproductive failure in infected individuals (B).

In addition to strongly reducing egg production, infection also reduced offspring viability. Across all infected females (n_Infected_ = 61), only six produced viable offspring, yielding a total of 14 nauplii. In contrast, unexposed females (n_Unexposed_ = 48) produced 1,976 nauplii in total, representing a >99% reduction in offspring production associated with infection. Notably, all infected females that successfully produced offspring were infected by sympatric parasites; however, given the small number of successful cases, this pattern should be interpreted cautiously.

### The timing of reproductive death

The timing of parasite-induced castration is a key determinant of its demographic consequences for host populations. Infected females produced viable offspring for a significantly shorter time window post-isolation compared to other treatments (GLMM_t3_, z = -9.54, p < 0.001). These effects emerged within days of infection (mean = 1.21 days ± 0.55 SE), indicating that interference from *S. solidus* is unlikely to arise solely from physical obstruction by growing parasites. Although infected females produced almost no viable offspring, many continued to generate additional clutches: 40 of 61 infected females produced at least one subsequent clutch, of which the majority failed to hatch (n = 35, 87.5%; Fig. 4B). This pattern underscores that castration is not due to an inability to produce eggs *per se*, but rather to a failure of eggs to survive or develop (see Fig. 3 for examples of non-viable eggs produced by females).

The timing and magnitude of this reproductive collapse in infected females is striking. Across all other treatments, females showed sustained clutch production over periods spanning weeks to months, consistent with long-term sperm storage in cyclopoid copepods (Fig. 4A). While infected females died significantly earlier than females in other treatments (GLMM_t4_, z = -2.86, p = 0.004; Fig. 2C), they typically survived well beyond their brief reproductive window, surviving on average for 45.5 days post-isolation (range = 7-103 days). This temporal disconnect between survival and reproduction suggests that *S. solidus* rapidly suppresses host fecundity long before causing host mortality, defining a prolonged period during which infected hosts remain alive but effectively sterile (existing as an extended parasite phenotype as a result of reproductive death). *S. solidus* infections at time of observation (9-14 days post exposure) were never cleared by copepods, aligning with previous research findings (van der Veen and Kurtz 2002).

### Fitness consequences of parasite resistance

Parasite exposure influenced female fecundity and offspring viability even in the absence of successful infection, but costs depended strongly on parasite origin. Females exposed to allopatric parasites produced significantly fewer eggs than both unexposed controls (GLMM_m1_, z = -2.22, p = 0.026) and sympatrically-exposed females (GLMM_m1_, z = -0.165, p = 0.0078; Fig. 4A), and this reduction translated into lower offspring viability: the proportion of viable nauplii was significantly reduced in allopatric-exposed females relative to controls (GLMM_m4_, z = - 2.07, p = 0.039; Fig. 4B).

In contrast, females exposed to sympatric parasites did not exhibit comparable reductions in fecundity or reproductive output. Egg production did not differ significantly between sympatric-exposed females and unexposed controls (GLMM_m2_, z = 1.02, p = 0.309), and sympatrically-exposed females exhibited significantly longer fecundity windows than all other treatments, with viable nauplii hatching up to 69 days post-isolation (GLMM_t3_, z = 2.09, p = 0.039). Together, these results show that exposure costs are real but genotype-dependent: allopatric exposure imposes measurable fecundity and viability penalties, while sympatric exposure appears to carry negligible cost and may even extend reproductive lifespan (Fig. 4).

### Mechanics underlying clutch production

To evaluate whether parasite interactions alter the timing of clutch production in gravid females, we analyzed intra-clutch holding time and egg output across treatments. In a sperm-storing species, distinguishing if females have fixed reproductive schedules versus being able to actively modulate clutch production provides critical insight to understanding how castration operates mechanistically.

We detected no significant treatment effect on the number of days females carry clutches (intra-clutch holding time, mean = 3.19 days), and the time between clutches did not differ between exposed and unexposed females (inter-clutch intervals, GLMM_t2_, z = 1.17, p = 0.242). Together, these results indicate that interacting with parasites does not disrupt the scheduling of clutch production.

Across all treatments, both eggs per clutch (GLMM_m1_, z = -7.88, p < 0.001) and nauplii survival per clutch (GLMM_m4_, z = -9.772, p < 0.001) declined significantly across successive clutches (Fig. 4C), consistent with progressive depletion of stored sperm or resources in isolated females. Clutch symmetry also decreased across clutch order, with egg production becoming increasingly uneven over time, further supporting cumulative reproductive constraint rather than active modulation of clutch composition (<u>Supp.</u> Fig. 1).

## Discussion

Parasite exposure is not a neutral event, even when infection fails. Hosts may either overcompensate (Minchella and Loverde 1981) or suppress fecundity (Horn and Luong 2018) as a result of evading infection. Successfully infective parasites (by definition) co-opt host resources, often reducing survival and reproductive output. These interactions shape both the dynamics of host populations and the genotypes of transmitted parasites. Despite this, we know little about the fitness consequences of parasite exposure in trophically transmitted systems. This gap is especially acute in first-intermediate hosts, where parasites are microscopic and their effects on host fitness are difficult to detect and rarely quantified (Hawley et al. 2011).

### Reciprocal asymmetries in parasite-host interactions

We considered diverse parasite origins, using both sympatric (Vancouver Island, Canada) and allopatric (Norway) host-parasite combinations; thus allowing comparison between host-parasite combinations with shared evolutionary history and those without recent ecological association.

We find that parasite exposure without successful infection can carry substantial fitness consequences, and that these effects depend strongly on parasite origin. Local adaptation is expected more often for highly virulent parasites, i.e. those that significantly increase host mortality or induce sterility, because they exert a stronger selective force on their hosts (Greischar and Koskella 2007). Indeed, infection by *S. solidus* was highly virulent, both significantly shortening survival and inducing castration in hosts. Sympatric parasites were more infective and achieved higher infection intensities (63% sympatric infection success versus 24% allopatric infection success, Fig. 2A), consistent with genotype-dependent variation in parasite performance observed in other host-parasite systems (Carius et al. 2001, Refardt and Ebert 2007, Kalbe et al. 2016). However, when infection was evaded, females exposed to sympatric parasites maintained high fecundity and survival. In contrast, allopatric parasites were less infective, but exposure was associated with reduced egg production, lower offspring viability, and shorter host lifespan.

These findings reveal a reciprocal asymmetry consistent with local adaptation: sympatric parasites are more infective but associated with lower costs when resisted, whereas allopatric parasites are less infective but associated with higher costs of exposure. Although only two parasite populations were examined, the magnitude and consistency of genotype-dependent differences suggest that parasite genotype plays a central role in shaping host fitness outcomes. Collectively, these results suggest that the selective consequences of host-parasite interactions extend well beyond successful infection. If exposure alone can reduce survival and reproduction, then the evolutionary dynamics of host-parasite systems may be governed as much by unsuccessful encounters as by successful ones. This reframes host-parasite interactions as a continuum of outcomes, where selection operates across all encounters, rather than being concentrated solely in infected hosts (Pfenning-Butterworth et al. 2023).

### Schistocephalus solidus as a castrating parasite in first intermediate hosts

Our results demonstrate that *S. solidus* functions as a true parasitic castrator in females of its first intermediate host, the cyclopoid copepod *Acanthocyclops robustus*. Infection by a single parasite resulted in rapid and near-complete suppression of host reproduction, often within 24 hours of parasite establishment. While parasitic castration is widespread across invertebrates, it typically emerges gradually with increasing parasite burden or developmental stage (Sorensen and Minchella 2001, Jensen et al. 2006, Best et al. 2010). The timing and dose-independent effects observed here distinguish the copepod-*S. solidus* interaction as an unusually rapid and virulent example of parasite-mediated castration.

The proximate mechanisms underlying castration in *S. solidus* remain unresolved. In other systems, parasitic castrators can disrupt host reproduction through physiological or endocrine pathways rather than simple resource depletion (Ebert et al. 2004, Hurd 2002, Lettini and Sukhdeo 2010). Infected females in our study produced subsequent clutches more rapidly than other treatments, but these clutches were rarely viable (Fig. 2), indicating that reproductive failure occurs downstream of egg production. This pattern is consistent with disruption of sperm storage or fertilization processes in females, although further work is needed to distinguish among mechanisms. Despite this uncertainty, the rapid cessation of reproduction suggests that *S. solidus* interferes directly with host reproductive regulation rather than acting solely through resource extraction. Such manipulation redirects host reproductive energy while maintaining host viability. This is consistent with theory predicting that castrators maximize fitness by suppressing reproduction without killing the host prematurely (O’Keefe and Antonovics 2002, Lafferty and Kuris 2009).

Sex-specific effects of infection further highlight the complexity of parasite impacts in this system. Most previous work on *S. solidus* in copepods has focused on males (Franz and Kurtz 2002, Kurtz and Hammerschmidt 2006, Benesh and Hafer 2012), likely because males are more susceptible to infection (Wedekind and Jakobsen 1998). Pilot studies (Fouilloux et al., *in prep*) demonstrate that infected males can successfully fertilize females, although the impact of infection in sperm quality and offspring performance remains unknown. Our results show that infection in females leads to severe reproductive suppression, emphasizing that the consequences of infection differ markedly between sexes. Accounting for these differences is essential for understanding how parasite effects scale to population-level dynamics, particularly in short-lived invertebrates where female reproductive output drives population growth (Hurd 2001).

### S. solidus as a trophically transmitted parasite

For trophically transmitted parasites, timing is everything. Parasites must keep intermediate hosts alive long enough to be predated, yet castrate them early enough to redirect reproductive energy toward parasite growth. Thus, the consequences of such a virulence-transmission tradeoff are high, where parasites that kill intermediate hosts prior to transmission incur complete fitness loss (O’Keefe and Antonovics 2001, Sorensen and Minchella 2001, Jensen et al. 2006). Here, we find that while infected copepods survive significantly shorter periods of time compared to unexposed and exposed treatments (similar to trematode infection in snails, Fredensborg et al. 2005 or bacterial infections in *Daphnia*: Ebert et al. 2004, Labbé et al. 2010, Gowler et al. 2023), they nevertheless can survive more than 40 days with their tapeworm parasite.

Although we did not monitor parasite survival explicitly, we did dissect dead females to confirm infection intensity and often observed that well developed, live parasites emerge from host bodies (see <u>Supp. Video 1</u>). Until now, we had little empirical data on how long infected first-intermediate hosts survive once infected with *S. solidus*. These data provide a critical baseline. With infection timing and host survival now quantified, we can begin to ask whether infected copepods fall within the predation window of sticklebacks in ecologically realistic contexts.

Infection by *S. solidus* is also known to affect fecundity in its second intermediate host, the threespine stickleback. The effect of *S. solidus* infection on fecundity in fish hosts varies across populations (increased/decreased clutch sizes, McPhail and Peacock 1983, Heins et al. 2010, Heins and Baker 2014) but none exhibit the rapid and complete reproductive suppression observed here in copepods. For a trophically transmitted parasite, this contrast is striking. Does *S. solidus* deploy the same physiological tactics across consecutive hosts, with outcomes differing due to invertebrate versus vertebrate immune architecture? Or does the parasite shift strategies entirely as it matures (Hébert et al. 2017)? These questions sit at the heart of how complex life cycles evolve.

Our results highlight that the earliest stages of infection represent a critical, and often overlooked, axis of selection. Exposure, resistance, and early parasite development act as a filter through which only a subset of parasite genotypes continue through the life cycle, while simultaneously shaping host population demography (and thus the future pool of susceptibles). In this view, transmission is not simply the outcome of successful infections, but the cumulative result of selection operating across all host-parasite encounters. Future work examining the mechanisms underlying castration, sex-specific responses, and host-specific strategies will further clarify how parasites navigate and exploit multi-host systems.

## Acknowledgements

This work was funded by the National Science Foundation EEID grant (Grant No. 2243076 to JLH and AH). A huge thank you to Jesse Weber for Norwegian *S. solidus* parasites and original copepod colonies and to Daniel Bolnick for helpful conversations. Also, a thank you to the beautiful copepods that bring endless fun and joy to our lives.

## Author contributions

CAF, JLH, and AH conceived the ideas and CAF designed methodology; CAF, JSC, IS, MLS, ALR, RP, and JS collected the data; CAF analyzed the data; CAF led the writing of the manuscript. All authors contributed critically to the drafts and gave final approval for publication. This work is a culmination of effort by researchers spanning undergraduate, graduate, to early- and mid-career stages, the diversity of which significantly contributed to the quality of this study.

## Data availability

All code, data, and figures are publicly available on <u>CAF’s GitHub</u>. These data will be submitted to the EDI repository upon manuscript acceptance.

