## SUPPLEMENTARY FIGURES for "From exposure to infection: divergent fitness consequences of parasite encounters in a trophically-transmitted system"

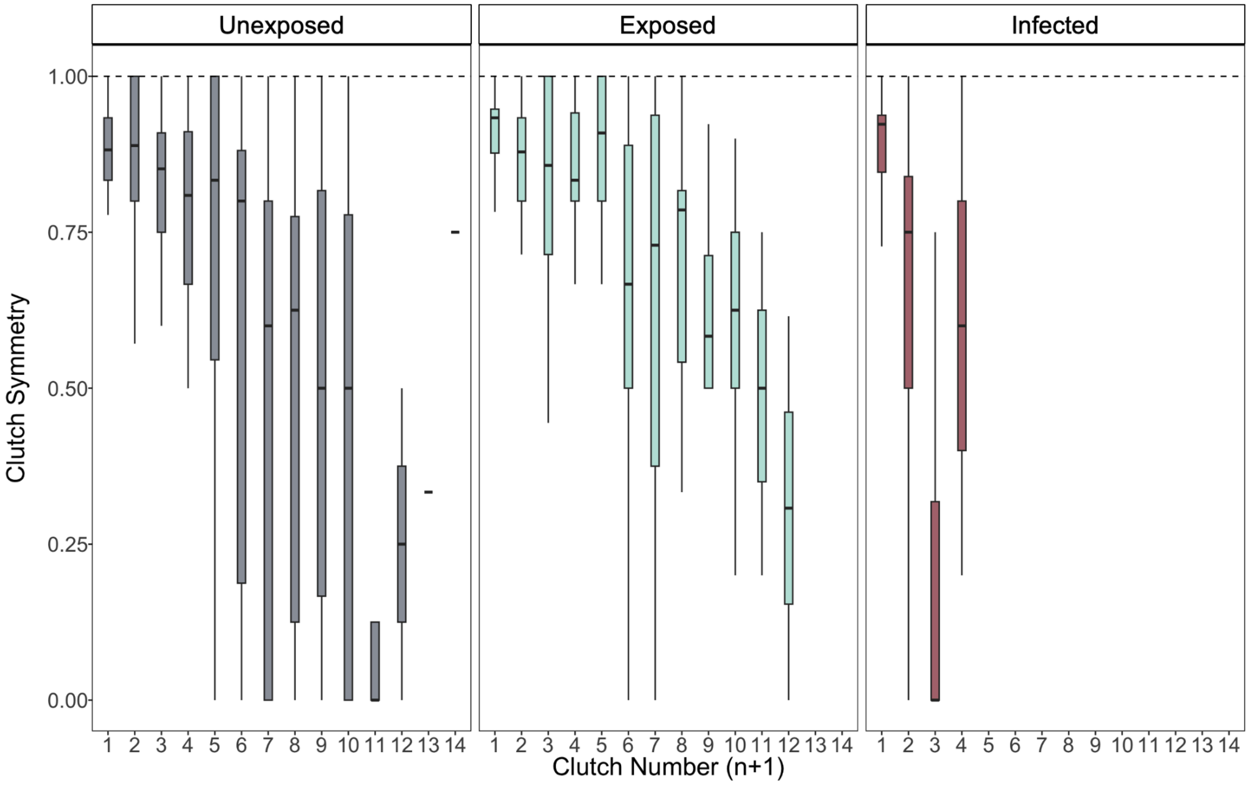


**Supplementary Figure 1.** Symmetry of clutches across sequential production in gravid females. Symmetry between clutches decreases across time. Dashed lines indicate perfect symmetry between left and right clutches. Boxplots denote median and 95 IQRs.

Clutch symmetry significantly decreases across clutch number ($z = -10.968$, $p < 0.001$, ordered beta glmmTMB). Larger clutches were less likely to show complete asymmetry (zi: $z = -7.02$, $p < 0.001$) and showed lower variance in symmetry (dispersion: $z = 7.90$, $p < 0.001$), suggesting that egg allocation becomes more consistent as clutch size increases. Infected and experimentally castrated females showed significantly higher variance in symmetry relative to controls (INF: $z = -3.76$, $p < 0.001$; EXP_Allopatric: $z = -3.01$, $p = 0.003$), indicating that treatment disrupts the consistency of egg distribution independently of mean symmetry.
